# Nuclear receptor HNF4A trans-represses CLOCK:BMAL1 and acts as a core component of tissue-specific circadian networks

**DOI:** 10.1101/424556

**Authors:** Meng Qu, Tomas Duffy, Tsuyoshi Hirota, Steve A. Kay

## Abstract

Either expression level or transcriptional activity of various nuclear receptors (NRs) have been demonstrated to be under circadian control. With a few exceptions, little is known about the roles of NRs as direct regulators of the circadian circuitry. Here we show that the nuclear receptor HNF4A strongly trans-represses the transcriptional activity of the CLOCK:BMAL1 heterodimer. We define a central role for HNF4A in maintaining cell-autonomous circadian oscillations in a tissue-specific manner in liver and colon cells. Not only transcript level but also genome-wide chromosome binding of HNF4A is rhythmically regulated in the mouse liver. ChIP-seq analyses revealed co-occupancy of HNF4A and CLOCK:BMAL1 at a wide array of metabolic genes involved in lipid, glucose and amino acid homeostasis. Taken together, we establish that HNF4A defines a novel feedback loop in tissue-specific mammalian oscillators and demonstrate its recruitment in the circadian regulation of metabolic pathways.

**Significance:** Interlocked feedback loops promote robustness and stability in a system and are a feature of circadian clocks in both animal and plants. The mammalian circadian clock is known to consist of two transcriptional feedback loops, relying on the transcriptional activity of the master complex CLOCK:BMAL1 and the feedback regulation by its target genes. Our research extends this knowledge by establishing a novel feedback loop in peripheral circadian oscillators and highlights the underlying mechanisms mediated by the unappreciated CLOCK:BMAL1 trans-repression activity of the circadian nuclear receptor HNF4A.

## Introduction

Circadian rhythms are intrinsic, entrainable oscillations of about 24 hours in behavior, metabolism and physiology found in almost all living organisms. This internal clock allows organisms to synchronize their physiology and behavior with the predictable cycle of day and night. Disruption of normal circadian rhythms leads to clinically relevant disorders including neurodegeneration, diabetes, obesity, and cardiovascular disease (1–4). In mammals, the core circadian circuit composed of transcriptional activators and repressors that form an autoregulatory transcriptional feedback loop, is necessary for the generation and regulation of circadian rhythms with an endogenous period close to 24 hr. The bHLH-PAS transcription factors CLOCK and BMAL1 are heterodimeric transcriptional activators that drive the expression of *Period* genes (*Per1, Per2*, and *Per3*) and *Cryptochrome* genes (*Cry1* and *Cry2*), by binding E-box cis-regulatory elements at promoters (5–9). Subsequently, the PER-CRY protein complex inhibits transcription of its own genes by directly inhibiting CLOCK:BMAL1 activity (8, 10–13). This feedback loop has been known as the critical hub of the mammalian oscillator.

While many mammalian peripheral tissues have circadian clocks, genes showing circadian expression are markedly diverse within individual tissues of mouse or human, indicating tissue-specific regulation of circadian output relevant to the function of the particular organ (14–17). The core clock may be regulated in a tissue-specific manner as well, supported by observations that explants of diverse mouse tissues express differences in circadian period and phase (16), and that output networks must be driven by the core clock components in a cell type-specific manner. However, such tissue-specific factors responsible for robust tissue-specific circadian rhythmicity remain to be identified.

The nuclear receptor (NR) superfamily is comprised of 48 members in human. NRs coordinate processes as diverse as development, reproduction, and many aspects of metabolism. As sensors for lipophilic vitamins, hormones, and dietary lipids, NRs canonically function as ligand-activated transcription factors that regulate the expression of their target genes to affect physiological pathways (18). The importance of NRs in maintaining optimal physiological homeostasis is illustrated in their identification as potential targets for therapeutic drug development to combat a diverse array of diseases, including reproductive disorders, inflammation, cancer, diabetes, cardiovascular disease, and obesity (19). Various NRs have been implicated as targets of the circadian clock, which may contribute to the circadian regulation of nutrient and energy metabolism. Over half of the NR family members are expressed in a rhythmic manner (20). Further, PER2 and CRYs were found to broadly interact with NRs and potentially modulate their transcriptional activity (21–24).

Apart from the circadian control of NR expression and transcriptional activity, other direct links between NR activity and circadian clock function have been identified via NRs’ role as circadian clock inputs. In addition to functions in development, metabolism and immunity, the nuclear receptors REV-ERBs and retinoic acid receptor-related orphan receptors (RORs) constitute the secondary feedback loop in the circadian cycle, by directly repressing and activating *Bmal1* expression, respectively, and in turn being transcriptionally regulated by CLOCK:BMAL1 (25–28). Dysregulation of REV-ERBs and RORs have dramatic effects on the robustness of circadian oscillators (29–33). However, little is known as to whether other NRs are involved in the regulation of core clock parameters.

The nuclear receptor hepatocyte nuclear factor 4A (HNF4A) binds as a homodimer to the consensus DNA sequence consisting of two direct repeats and controls the expression of downstream genes (34). It is expressed in a tissue-specific manner and plays essential roles in the development and physiology of liver, pancreas, kidney and intestines, particularly lipid and glucose metabolism and inflammation (35–41). In the current study, we demonstrate that HNF4A potently inhibits the transcriptional activity of CLOCK:BMAL1 heterodimer. Furthermore, our data clearly show that HNF4A is necessary for the regulation of intrinsic circadian oscillations in liver and colon cells, suggesting it to be a novel essential component of tissue-specific clock networks.

## Results

### Identification of HNF4A as a CLOCK:BMAL1 trans-repressor

The growing body of evidence demonstrating a link between nuclear receptors, metabolic pathways and intrinsic circadian rhythmicity (20–24) prompted us to test the potential for a broader range of nuclear receptors in mediating this crosstalk. In a physical interaction profiling, we noted a robust binding between core clock proteins and the HNF4A protein, as shown in co-immunoprecipitation experiments using either recombinant or endogenous proteins (Fig. ***1A-C***). To evaluate the biological significance of the interactions, we hypothesized that either the core clock components regulate the activity of nuclear receptor HNF4A, or HNF4A may transduce metabolic signals as inputs to the circadian system. To test the first hypothesis, we conducted a reporter assay in HEK 293T cells using a luciferase reporter driven by tandem HNF4A consensus binding motifs. Neither CRY1 nor CLOCK:BMAL1 co-expression affected the HNF4A transcriptional activity, indicating that HNF4A activity is unlikely to be directly regulated by these clock proteins (Fig. ***S1***).

**Fig. 1.**
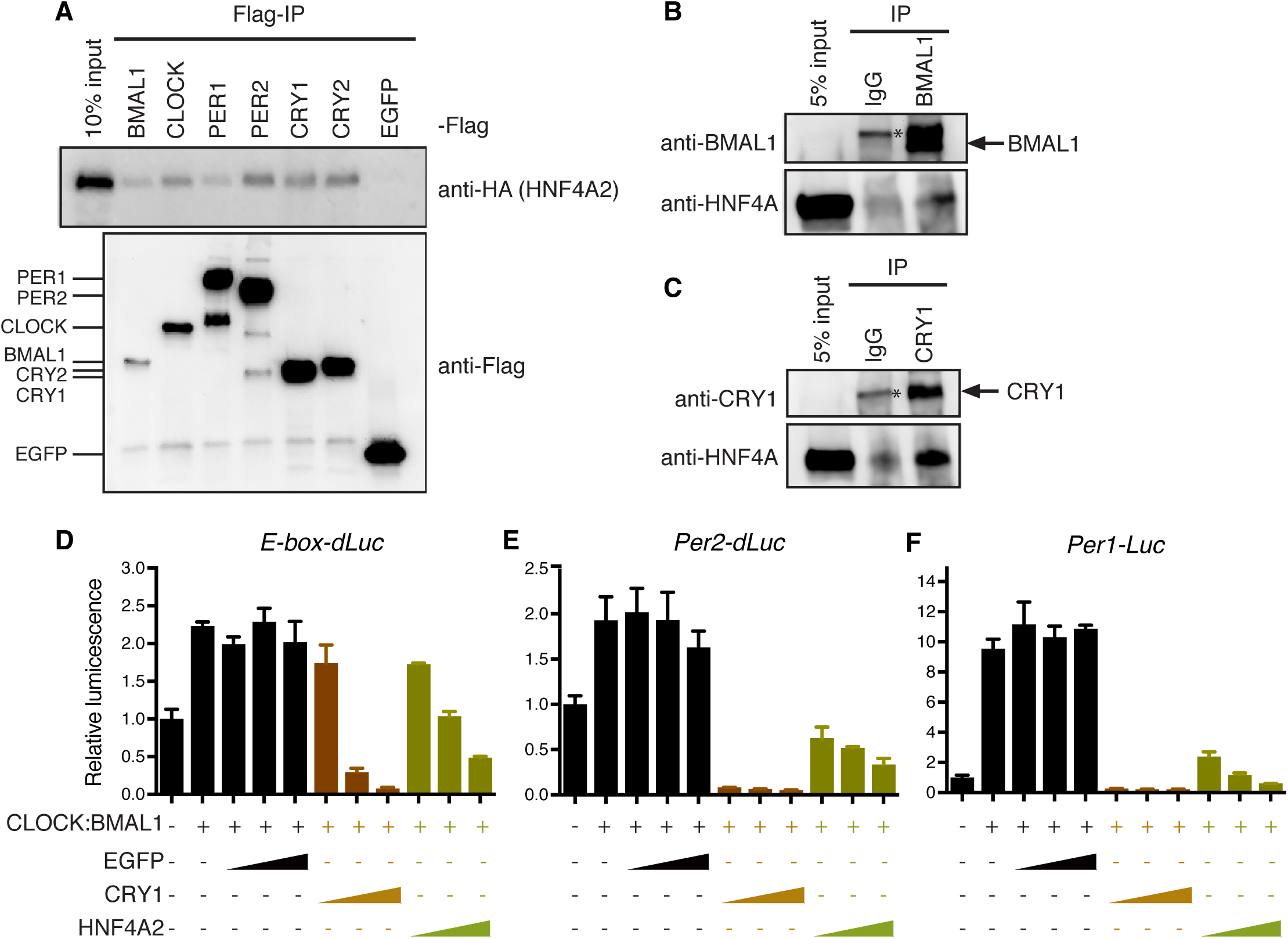
HNF4A physically interacts with the core clock complex and trans-suppresses the transcriptional activity of CLOCK:BMAL1. (A) Co-immunoprecipitation of transiently expressed core clock proteins and HNF4A from HEK 293T extract. (B-C) Co-immunoprecipitation of endogenous BMAL1 (B) or CRY1 (C), and HNF4A from HepG2 extract. (D-F) HNF4A inhibits CLOCK:BMAL1 activity in luminescent reporter assays. HEK 293T cells were co-transfected with reporter gene *E-box-dLuc* (D), *Per2-dLuc* (E), or *Per1-Luc* (F), CLOCK:BMAL1, and increasing amounts of EGFP, CRY1, or HNF4A plasmid (n = 3 for each condition, mean ± SD).

We then tested the second hypothesis, asking whether the core clock feedback loops are regulated by the HNF4A protein. As HNF4A consistently binds CLOCK:BMAL1, we evaluated the effect of HNF4A on CLOCK:BMAL1 activity in endpoint reporter assays using the *E-box-dLuc, Per1-Luc* or *Per2-dLuc* luciferase reporters. Surprisingly, we observed a consistent inhibition of CLOCK:BMAL1 activity by HNF4A co-expression in a dose-dependent pattern, similar to the canonical clock repressor CRY1 (Fig. ***1D-F***). Collectively, these findings indicate a novel activity of the nuclear receptor HNF4A in trans-repressing another transcription factor CLOCK:BMAL1 and prompted us to evaluate its role in the intact molecular circadian oscillators.

### HNF4A is essential for tissue-specific rhythm maintenance and period regulation

As HNF4A is minimally expressed in bone osteosarcoma U2OS cells, our previous genome-wide RNAi screen (42) was unable to assess its potential as a clock modifier (Fig. ***S2***). We therefore screened cell lines isolated from various tissue types to identify ones that both express a high level of HNF4A and oscillate upon dexamethasone synchronization. The cell lines we eventually identified are human liver cells Hep3B, mouse liver cells dihXY and human colon cells SW480, which were genetically modified to stably express *Per2-dLuc* or *Bmal1-Luc* reporter gene. To assess the role of HNF4A in regulating circadian oscillations of these cells, we performed gene knockdown by using *Hnf4a* or *Cry1* siRNAs selected for their potency (42) (Fig. ***S3A-C***). Surprisingly, the effect of *Hnf4a* knockdown in both liver cell lines Hep3B and dihXY was even more disruptive than *Cry1* knockdown, leading to arrhythmicity (Fig. ***2A-D***). In SW480 cells, in contrast, the effect of *Hnf4a* knockdown on the circadian rhythm was milder and resulted in period shortening by roughly one hour (Fig. ***2E*** **and** ***F***), similar to the *Cry1* knockdown (Fig. ***S3D* and *E***). The cell type-specific phenotypes of *Hnf4a* knockdown are consistent with previous observations for other clock genes (43).

**Fig. 2.**
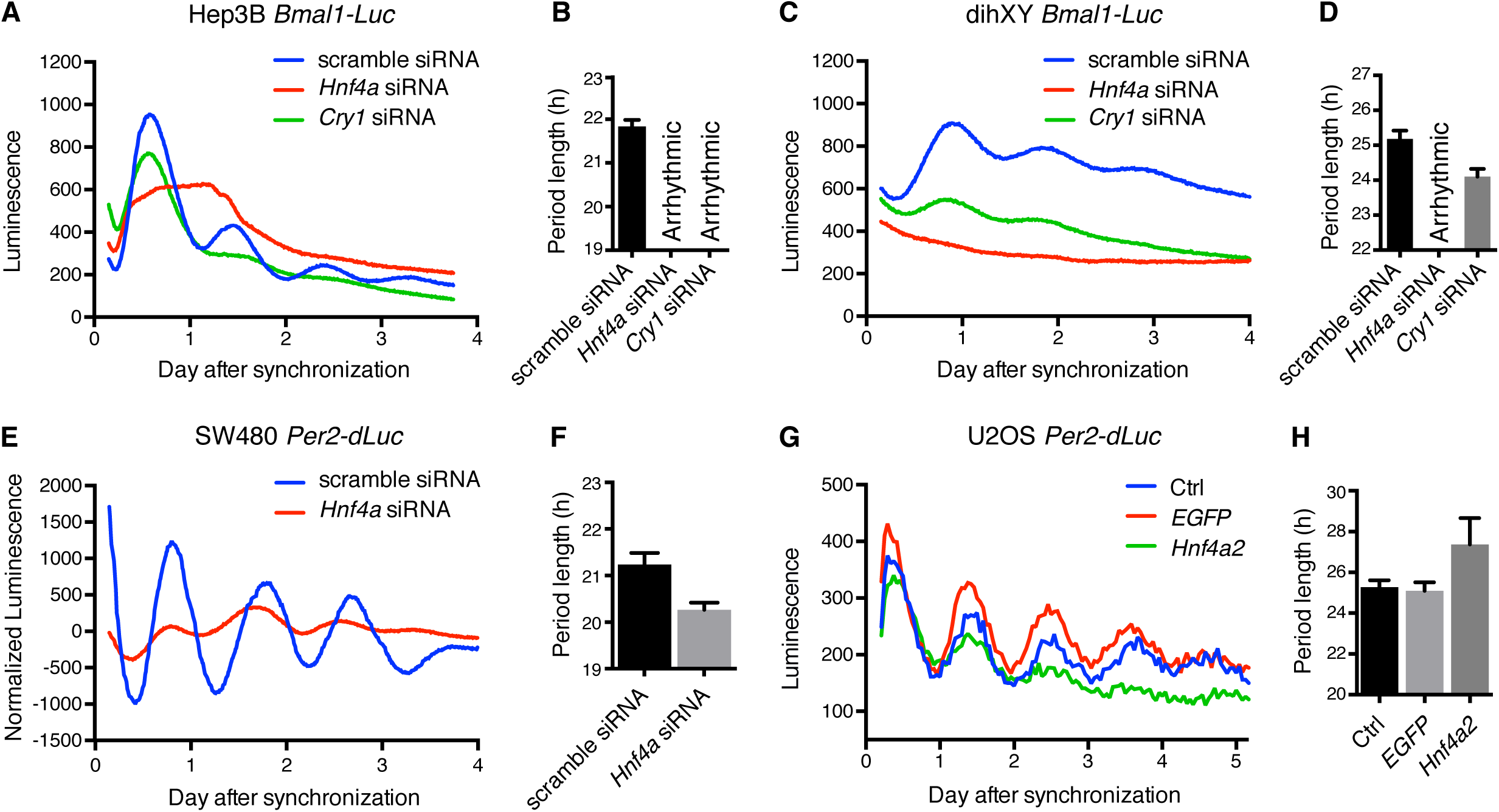
HNF4A is critical for circadian rhythm maintenance and period regulation in liver and colon cells. (A) Effect of control scramble, *Cry1*, or *Hnf4a* siRNA on *Bmal1-Luc* reporter in human liver cells Hep3B (n = 5). (B) Mean period of circadian luminescence rhythms in (A). Data are represented as mean ± SD (n = 5). (C) Effect of control scramble, *Cry1*, or *Hnf4a* siRNA on *Bmal1-Luc* reporter in mouse liver cells dihXY (n = 5). (D) Mean period of circadian luminescence rhythms in (C). Data are represented as mean ± SD (n = 5). Significance between scramble siRNA and *Cry1* siRNA was assessed by Student’s t test, p= 7.75e-05 (< 0.05). (E) Effect of control scramble or *Hnf4a* siRNA on *Per2-dLuc* reporter in human colon cells SW480 (n = 3). (F) Mean period of circadian luminescence rhythms in (E). Data are represented as mean ± SD (n = 3). Significance was assessed by Student’s t test, p=4.72e-03 (< 0.05). (G) Effect of expressing EGFP or HNF4A on *Per2-dLuc* reporter in U2OS cells. (H) Mean period of circadian luminescence rhythms in (G). Data are represented as mean ± SD (n = 16). Significance between EGFP and HNF4A expression was assessed by Student’s t test, p= 1.91e-05 (< 0.05).

Using U2OS *Per2-dLuc* cells, we generated stable lines expressing EGFP, CRY1, PER2, or HNF4A. Additional expression of CRY1 and PER2 proteins led to arrhythmicity along with greatly reduced *Per2-dLuc* reporter signals (Fig. ***S3F***). In contrast, ectopic expression of the *Hnf4a* gene significantly dampened the amplitude and lengthened the period (Fig. ***2G*** **and** ***H***). Taken together, our data suggest that HNF4A is essential for circadian rhythmicity of cells where it is endogenously expressed, and significantly alters the intrinsic circadian clock when ectopically expressed. We thus identify HNF4A to be a central regulator of tissue-specific circadian rhythms in liver and colon cells.

### HNF4A acts differently from other CLOCK:BMAL1 repressors

As HNF4A is able to bind CRY proteins as well (Fig. ***1A*** **and** ***C***), we asked whether its CLOCK:BMAL1 repression is attributable to an indirect mechanism bridged by CRYs. We applied the CLOCK HI:BMAL1 LL/AA mutant which can no longer be sequestered by CRY1 (44, 45). Even though the activation of *Per1-Luc* reporter by CLOCK HI:BMAL1 LL/AA mutant is less pronounced than the wild-type complex, our results show that CRY1 repression was substantially abolished (Fig. ***3A***). In contrast, HNF4A could still prominently suppress the mutant complex, suggesting that HNF4A inhibits CLOCK:BMAL1 by a mechanism independent of CRY1 (Fig. ***3B***). This conclusion was further supported by our observation that the knockdown of two *Cry* genes (42) did not affect the extent of HNF4A-dependent repression of CLOCK:BMAL1 activity (Fig. ***S4A***).

**Fig. 3.**
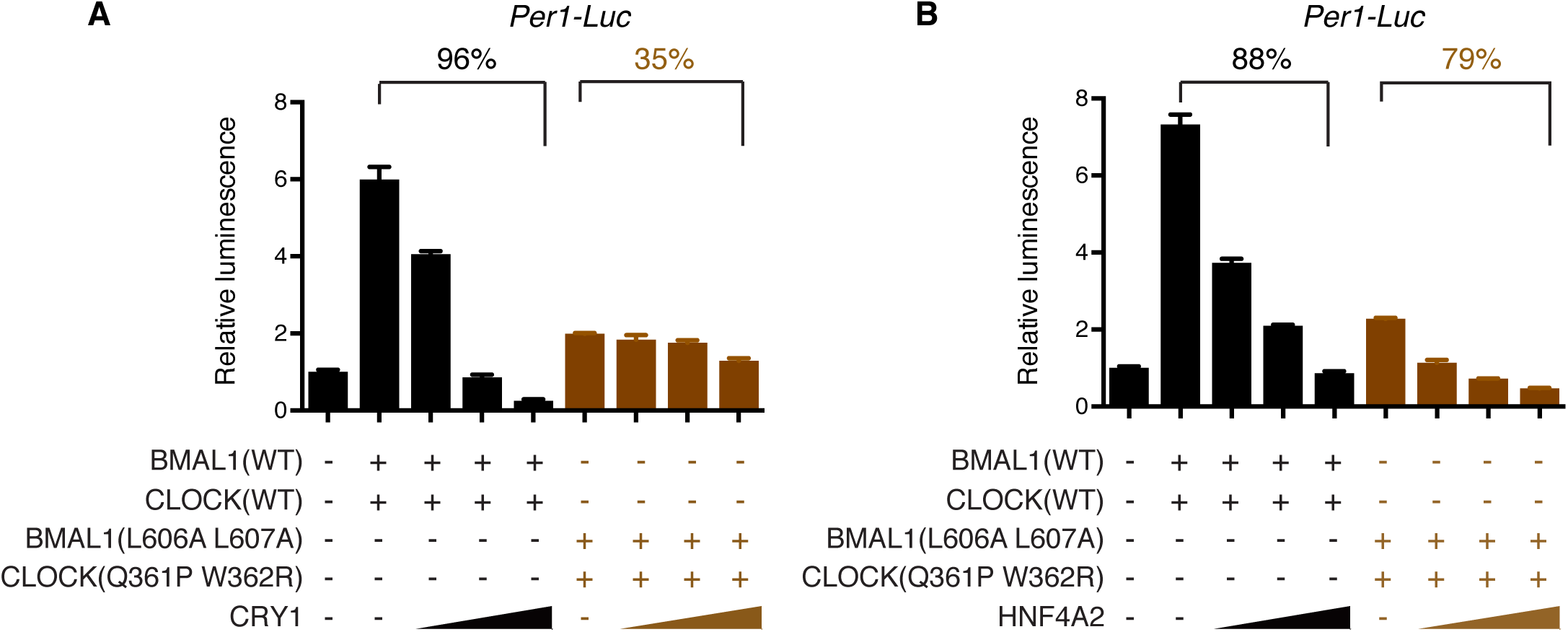
HNF4A targets CLOCK:BMAL1 differently from CRY1. (A-B) HNF4A is able to suppress a CLOCK:BMAL1 mutant which interferes with sequestration by CRY1. HEK 293T cells were co-transfected with reporter gene *Per1-Luc*, wild type or mutant CLOCK:BMAL1, and increasing amounts of CRY1 (A) or HNF4A (B) plasmid (n = 3 for each condition, mean ± SD). Percentages on the top indicate the degree of CLOCK:BMAL1 repression.

Cancer/testis antigen PASD1 was also identified to be able to repress the CLOCK:BMAL1 activity, dependent upon exon 19 of the *Clock* gene (46). Deletion of exon 19 was found to attenuate the transactivation potential of the transcription factor (6, 7). We found that HNF4A could still potently repress the residual activity of the CLOCK^Δ19^:BMAL1 complex (Fig. ***S4B***). Consequently, there appears to be a novel targeting mechanism leveraged by HNF4A in achieving CLOCK:BMAL1 repression activity.

### Multiple HNF4A domains are involved in CLOCK:BMAL1 repression

Like most other nuclear receptors, HNF4A contains six structural domains that are necessary to mediate disparate functions: an N-terminal activation function (AF)-1 domain; a DNA-binding domain (DBD); a putative ligand-binding domain (LBD); a flexible hinge between the DBD and LBD; a C-terminal activation domain (AF-2), and a repressor region (F domain) that recruits corepressors and inhibits access of coactivators to AF-2 (47–52). We used various genetic tools to characterize the novel trans-repression activity of HNF4A. The predominant HNF4A splice variants, HNF4A1, HNF4A2 and HNF4A8, driven by tissue-specific promoters P1 or P2, are considered to have different physiological roles in development and transcriptional regulation of target genes. P1-driven HNF4A1 and HNF4A2, which differ exclusively by 10 amino acids in the F domain, are specifically expressed in the adult liver and kidney; whereas the P2-driven HNF4A8 is not found in these tissues but in fetal liver and the adult pancreas. Relative to HNF4A1/2, HNF4A8 lacks the N-terminal activation domain AF-1 (53–55). We further tested the effect of HNF4G, which shares nearly identical DBD and LBD domains with HNF4A. Intriguingly, all HNF4A isoforms we tested, together with HNF4G, potently repressed CLOCK:BMAL1 activity (Fig. ***4A***). It suggests that the HNF4 proteins share a common trans-repression activity in various peripheral tissues where they are specifically expressed, and the unshared structural domains or protein sequences are dispensable for the activity. To explore contribution of some major functional domains, we generated constructs expressing HNF4A2 fragments. We found that deletion of the AF-1 and DBD domains (Δ 1-119), LBD (Δ 137-365), or the F domain (aa 1-376) did not affect the protein expression (Fig. ***S5***) but substantially abolished an efficient repression of the CLOCK:BMAL1 activity (Fig. ***4B***). To potently trans-repress the CLOCK:BMAL1 transcriptional activity, our data reflect requirements of the DBD, LBD and F domains within the intact HNF4 protein.

**Fig. 4.**
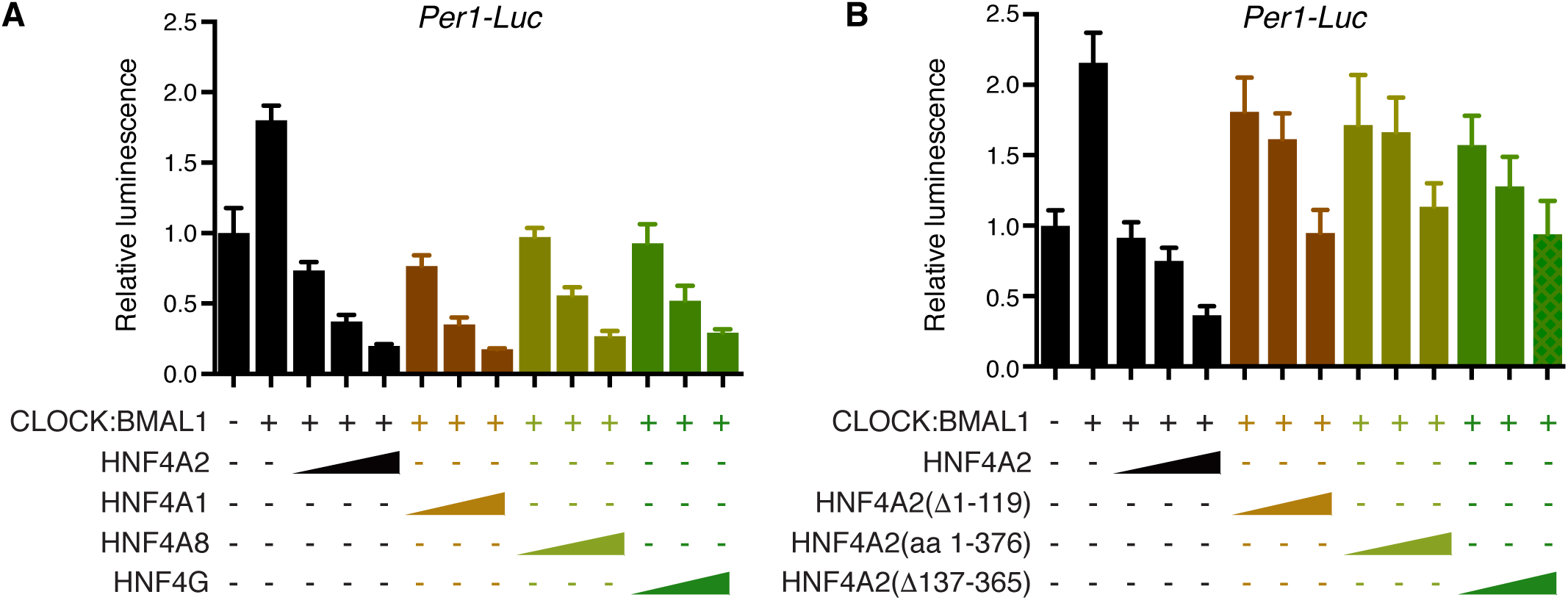
Multiple HNF4A functional domains are required for CLOCK:BMAL1 trans-repression. (A) HNF4A isoforms and HNF4G repress CLOCK:BMAL1 activity similarly. HEK 293T cells were co-transfected with reporter gene *Per1-Luc*, CLOCK:BMAL1, and increasing amounts of HNF4 plasmid (n = 3 for each condition, mean ± SD). (B) Domain analysis of HNF4A2. HEK 293T cells were co-transfected with reporter gene *Per1-Luc*, CLOCK:BMAL1, and increasing amounts of HNF4A2 fragment plasmid (n = 3 for each condition, mean ± SD).

### HNF4A mRNA level and chromatin binding are rhythmic in mouse

Our combined data suggest that HNF4A shares many of the canonical properties of a core clock protein (56). We further examined *Hnf4a* mRNA expression in liver, pancreas and colon of C57BL/6J mice over an LD cycle, and found it to be rhythmic on the circadian timescale, with a cyclic pattern (peak at night) and modest but detectable amplitude comparable to the *Cry2* gene (SAS PROC ANOVA, p<0.001, Fig. ***5***). HNF4A has been known to play central roles in dietary lipid absorbance, bile acid synthesis, conjugation, and transport (37, 57–59). However, the accumulation of HNF4A transcripts in the early evening is unlikely to be induced by diet, as mice that are normally fed and fasted have similar levels of *Hnf4a* transcript at ZT14, in contrast to the remarkable differences in gluconeogenic genes *Pck1* and *G6pc* (Fig. ***S6***). By analyzing ChIP-seq data (60), we observed binding of CLOCK, BMAL1 and CRYs at the *Hnf4a* gene (Fig. ***S7***), suggesting that *Hnf4a* transcription is directly modulated by the core clock proteins.

**Fig. 5.**
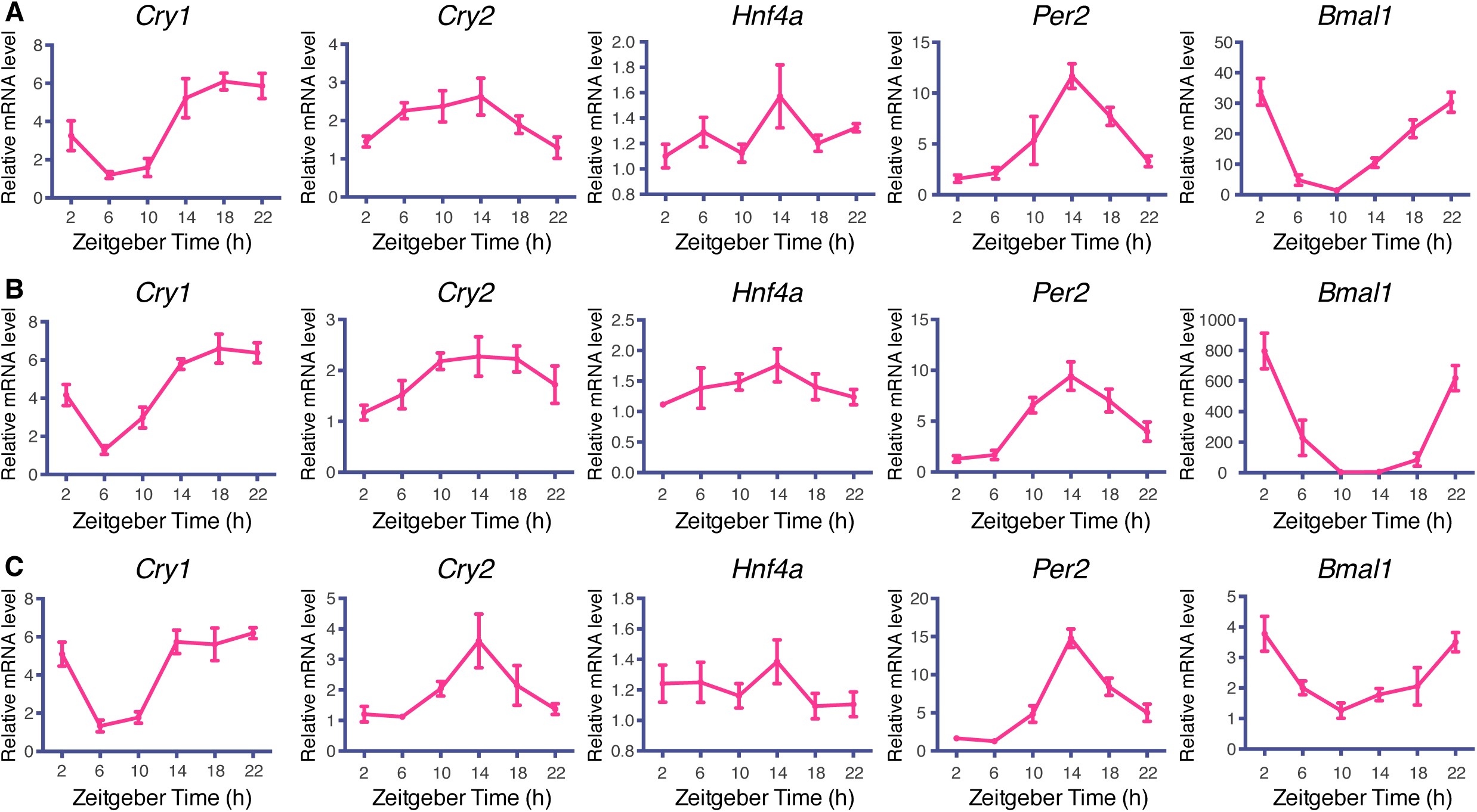
Gene *Hnf4a* is rhythmically expressed in vivo. Mouse liver (A), pancreas (B) and colon (C) were harvested at 4-h intervals over the course of 24 h. Transcript levels were analyzed by using RT-qPCR. Displayed are the means ± SD (n=5 mice) normalized to non-oscillating *Rplp0* expression levels.

In order to address whether the HNF4A functions are also rhythmically regulated, we used chromatin immunoprecipitation to evaluate its genome-wide binding at ZT4 and ZT16, which correspond to the trough and peak of *Hnf4a* transcript abundance. Surprisingly, out of 20,230 sites identified at ZT16, 17,839 are differentially occupied by HNF4A at ZT4, when only 6,163 peak sites were identified using the same cutoff (Fig. ***6A***). Consistently, genome-wide HNF4A occupancy is clearly more extensive at ZT16 compared to ZT4 (Fig. ***6B***). In particular, we observed strong rhythmicity of HNF4A binding at genes involved in well-characterized HNF4A-regulated processes (Fig. ***S8***). HNF4A induces the expression of *Hnf1a*, another tissue-specific transcription factor that plays essential roles in key developmental processes (61). HNF4A is also required for bile acid synthesis by directly targeting *Cyp7a1* (59, 62) and is required for the expression of the xenobiotic nuclear receptor PXR (NR1I2) (63). Indeed, all the three target genes show oscillating expression in the mouse liver with phases similar to *Hnf4a*, by exhibiting higher levels during the night (Fig. ***S9***, data derived from the CircaDB database) (64). In aggregate, our data strongly suggests that the oscillation of tissue-specific nuclear receptor HNF4A directly determines the rhythmicity of an array of its target genes, such that temporal regulation of various aspects of physiology can be achieved.

**Fig. 6.**
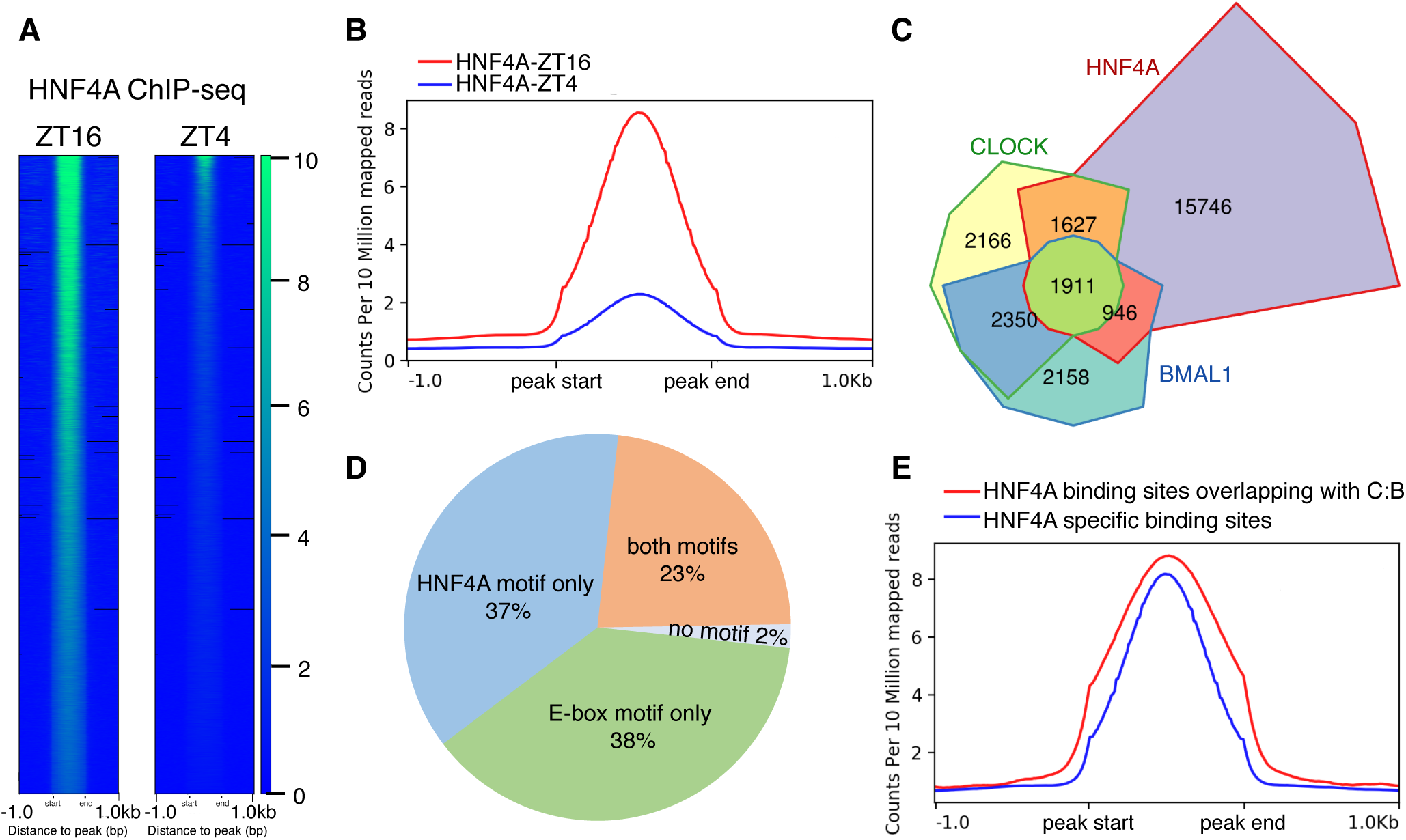
The genome-wide chromatin binding of HNF4A is rhythmic and broadly overlaps with that of CLOCK:BMAL1. (A) Heatmap views of genome-wide binding of HNF4A at ZT4 and ZT16, measured over 2 kb fragments flanking each peak. Peaks are ordered vertically by signal strength, represented by the green-blue gradient. (B) Meta-plot comparing genome-wide HNF4A binding intensity at ZT4 and ZT16. (C) Chow-Ruskey diagram showing genome-wide co-occupancy of BMAL1, CLOCK and HNF4A. An overlap was called when peaks overlap by >1bp. The boundaries for each protein are color-coded: BMAL1 (blue), CLOCK (green), and HNF4A (red). The areas of each domain are proportional to the number of binding sites. (D) Pie chart showing the distribution of E-box and HNF4A consensus motifs at BMAL1-CLOCK-HNF4A co-occupied peaks. (E) Meta-plot comparing HNF4A binding intensity at loci with and without CLOCK:BMAL1 co-occupancy.

### HNF4A and CLOCK:BMAL1 co-occupy clock and metabolic genes

To assess the genome-wide effects of HNF4A-mediated CLOCK:BMAL1 trans-repression, we compared cistromes of HNF4A (ZT16), BMAL1, and CLOCK (60) and calculated their overlaps. At the center of the Chow-Ruskey plot, 1911 loci in the genome are bound by all three factors (Fig. ***6C***). Motif analysis of the common binding sites shows a significant enrichment for the canonical CLOCK:BMAL1 E-box motif (CACGTG) and HNF4A-binding motif (Fig. ***S10A***). Interestingly, at 75% of the co-occupied peaks only one of the two most abundant motifs was identified (Fig. ***6D***). Furthermore, the HNF4A binding was stronger at loci co-occupied by CLOCK:BMAL1 relative to the HNF4A-exclusive binding sites (Fig. ***6E***). These results indicate that HNF4A and CLOCK:BMAL1 broadly interact at their co-occupied genomic loci, and a tethering mechanism may be involved in their co-regulation of target gene transcription. Gene Ontology analysis of genes commonly occupied by the two transcription factors shows enrichment for circadian circuits, as well as pathways regulating lipid, glucose and amino acid metabolism (Fig. ***S10B***). Hence, apart from the circadian rhythms, HNF4A may modulate the core metabolic processes by antagonizing CLOCK:BMAL1-facilitated transcriptional activation.

## Discussion

Our findings in this study highlight the fundamental roles played by the nuclear receptor HNF4s (including HNF4A and HNF4G) in maintaining the tissue-specific circadian oscillations in the intestinal tract and liver. Similarly, the *Bmal1* activator RORγ, also a nuclear receptor, controls the amplitude of clock genes in peripheral tissues without affecting rhythms in the SCN, where it is not expressed (28, 32). The identification of tissue-specific clock regulators reveals one mechanism generating tissue-specific circadian rhythms and allows precise modulation of clock networks in specific organs.

We report here a noncanonical trans-repression activity of HNF4A reminiscent of the anti-inflammatory activities of glucocorticoid receptor (GR) that are caused predominantly by regulating the key pro-inflammatory transcription factors nuclear factor-κB (NF-κB) and activator protein 1 (AP-1). Consistent with our observations here, nuclear receptor GR physically interacts with NF-κB or AP-1 heterodimers and represses their transcriptional activity (65, 66). The exact mechanism by which GR inhibits the pro-inflammatory factors is still unclear, although several models have been proposed. The chromatin recruitment of GR to the vicinity of the pro-inflammatory factors could be achieved by “tethering”, through the protein-protein interactions. Apart from this, at some promoters, it could also be mediated by a “composite binding” in which GR binding to both the target transcription factor and DNA is required. Afterwards, the first mode of GR-mediated trans-repression is attributable to a direct repression of the DNA binding and/or transactivation functions of the pro-inflammatory factors, dependent on the specific physical interactions. In addition, there are indirect modes that may involve competition for limiting amounts of coactivator, recruitment of corepressor, and interference with the basal transcriptional machinery (BTM) (67–69).

Our deletion analysis suggests involvement of the DBD, LBD and F domains of HNF4A in the trans-repression of CLOCK:BMAL1. Similarly, GR DBD and LBD domains are also demonstrated to be required for its trans-repression function (70–72). In HNF4A, the C-terminal F domain is known as a repressor region that is responsible for recruitment of corepressor SMRT/NCOR2. The requirement of the F domain may imply a role for corepressor recruitment in the process of CLOCK:BMAL1 inhibition. Taken together, we propose a novel activity of HNF4A in trans-repressing CLOCK:BMAL1, the core positive arm of the molecular clock, and consequently a central role in the maintenance of circadian oscillations. Supporting this model, we observed co-occupancy of HNF4A and CLOCK:BMAL1 at clock genes *Dbp, Cry1, Cry2, Per2* and *Per3* (Fig. ***S11***). An HNF4-binding motif was identified at most of these co-occupied loci, suggesting that throughout the HNF4A-mediated CLOCK:BMAL1 trans-repression at these clock genes, the “composite binding” mode may be at play. Hence, similar to GR, HNF4A may bind to the regulatory regions of the clock genes, trans-repressing the co-occupying CLOCK:BMAL1 via physical interactions.

Our findings also suggest HNF4A to be a central regulator in the circadian control of metabolic pathways. This role could be accomplished through HNF4A activities at two distinct levels, depending on the chromatin environment. First, the extensive co-occupancy of HNF4A and CLOCK:BMAL1 strongly suggest that, apart from the core clock, HNF4A negatively modulates the transcriptional activity of CLOCK:BMAL1 in diverse metabolic pathways (73–76). Further, the circadian expression and genome binding of the nuclear receptor HNF4A itself, particularly at loci not co-occupied by CLOCK:BMAL1, is implied to directly result in the circadian expression of downstream metabolic genes responsible for bile acid synthesis and xenobiotic metabolism. Taken together, HNF4A provides an interesting link between the molecular regulation of core clock proteins and the metabolic pathways, a subject for future studies.

## Materials and Methods

For details on luciferase assays, circadian assays, co-IP, qPCR, and HNF4A ChIP-seq and data analysis, please see *SI Materials and Methods*.

All animal care and experiments were performed under the institutional protocols approved by the Institutional Animal Care and Use Committee (IACUC) at USC.

## Acknowledgement

We thank Carrie Partch at UCSC for sharing SW480 *Per2-dluc* reporter cells; Michael Karin and Jeremy Rich at UCSD for sharing Hep3B *Bmal1-luc*, dihXY *Bmal1-luc* reporter cells. We appreciate Han Qu at UC Riverside for assisting statistical analysis. We also thank the NGS and Microarray Core Facility at TSRI for processing HNF4A ChIP-seq. This work was supported by National Institute of Diabetes and Digestive and Kidney Diseases grant 5R01DK108087 to SAK.

## Author Contributions

M.Q. and S.A.K. designed the research. M.Q. performed and analyzed the experiments. T.D. and M.Q. performed the bioinformatic analysis. M.Q., T.H. and S.A.K. wrote the manuscript.

## References

1. Chen L, Yang G (2015) Recent advances in circadian rhythms in cardiovascular system. Front Pharmacol 6. doi:10.3389/fphar.2015.00071.

2. Hastings MH, Goedert M (2013) Circadian clocks and neurodegenerative diseases: time to aggregate? Curr Opin Neurobiol 23(5):880–887.

3. Marcheva B, et al. (2010) Disruption of the clock components CLOCK and BMAL1 leads to hypoinsulinaemia and diabetes. Nature 466(7306):627–631.

4. Turek FW, et al. (2005) Obesity and Metabolic Syndrome in Circadian Clock Mutant Mice. Science 308(5724):1043–1045.

5. Bunger MK, et al. (2000) Mop3 Is an Essential Component of the Master Circadian Pacemaker in Mammals. Cell 103(7):1009–1017.

6. Gekakis N, et al. (1998) Role of the CLOCK Protein in the Mammalian Circadian Mechanism. Science 280(5369):1564–1569.

7. King DP, et al. (1997) Positional Cloning of the Mouse Circadian ClockGene. Cell 89(4):641–653.

8. Kume K, et al. (1999) mCRY1 and mCRY2 Are Essential Components of the Negative Limb of the Circadian Clock Feedback Loop. Cell 98(2):193–205.

9. Zheng B, et al. (2001) Nonredundant Roles of the mPer1 and mPer2 Genes in the Mammalian Circadian Clock. Cell 105(5):683–694.

10. Griffin EA, Staknis D, Weitz CJ (1999) Light-Independent Role of CRY1 and CRY2 in the Mammalian Circadian Clock. Science 286(5440):768–771.

11. Lee C, Etchegaray J-P, Cagampang FRA, Loudon ASI, Reppert SM (2001) Posttranslational Mechanisms Regulate the Mammalian Circadian Clock. Cell 107(7):855–867.

12. Sato TK, et al. (2006) Feedback repression is required for mammalian circadian clock function. Nat Genet 38(3):312–319.

13. Rosensweig C, et al. (2018) An evolutionary hotspot defines functional differences between CRYPTOCHROMES. Nat Commun 9(1):1138.

14. Panda S, et al. (2002) Coordinated Transcription of Key Pathways in the Mouse by the Circadian Clock. Cell 109(3):307–320.

15. Storch K-F, et al. (2002) Extensive and divergent circadian gene expression in liver and heart. Nature 417(6884):78–83.

16. Yoo S-H, et al. (2004) PERIOD2::LUCIFERASE real-time reporting of circadian dynamics reveals persistent circadian oscillations in mouse peripheral tissues. Proc Natl Acad Sci 101(15):5339–5346.

17. Ruben MD, et al. (2018) A database of tissue-specific rhythmically expressed human genes has potential applications in circadian medicine. Sci Transl Med:8.

18. Evans RM, Mangelsdorf DJ (2014) Nuclear Receptors, RXR, and the Big Bang. Cell 157(1):255–266.

19. Chawla A, Repa JJ, Evans RM, Mangelsdorf DJ (2001) Nuclear Receptors and Lipid Physiology: Opening the X-Files. Science 294(5548):1866–1870.

20. Yang X, et al. (2006) Nuclear Receptor Expression Links the Circadian Clock to Metabolism. Cell 126(4):801–810.

21. Han D-H, Lee Y-J, Kim K, Kim C-J, Cho S (2014) Modulation of glucocorticoid receptor induction properties by core circadian clock proteins. Mol Cell Endocrinol 383(1):170–180.

22. Kriebs A, et al. (2017) Circadian repressors CRY1 and CRY2 broadly interact with nuclear receptors and modulate transcriptional activity. Proc Natl Acad Sci 114(33):8776–8781.

23. Lamia KA, et al. (2011) Cryptochromes mediate rhythmic repression of the glucocorticoid receptor. Nature 480(7378):552–556.

24. Schmutz I, Ripperger JA, Baeriswyl-Aebischer S, Albrecht U (2010) The mammalian clock component PERIOD2 coordinates circadian output by interaction with nuclear receptors. Genes Dev 24(4):345–357.

25. Akashi M, Takumi T (2005) The orphan nuclear receptor RORα regulates circadian transcription of the mammalian core-clock Bmal1. Nat Struct Mol Biol 12(5):441–448.

26. Kim YH, et al. (2018) Rev-erbα dynamically modulates chromatin looping to control circadian gene transcription. Science 359(6381):1274–1277.

27. Preitner N, et al. (2002) The Orphan Nuclear Receptor REV-ERBα Controls Circadian Transcription within the Positive Limb of the Mammalian Circadian Oscillator. Cell 110(2):251–260.

28. Sato TK, et al. (2004) A Functional Genomics Strategy Reveals Rora as a Component of the Mammalian Circadian Clock. Neuron 43(4):527–537.

29. Bugge A, et al. (2012) Rev-erbα and Rev-erbβ coordinately protect the circadian clock and normal metabolic function. Genes Dev 26(7):657–667.

30. Cho H, et al. (2012) Regulation of circadian behaviour and metabolism by REV-ERB-α and REV-ERB-β. Nature 485(7396):123–127.

31. Liu AC, et al. (2008) Redundant Function of REV-ERBα and β and Non-Essential Role for Bmal1 Cycling in Transcriptional Regulation of Intracellular Circadian Rhythms. PLoS Genet 4(2):e1000023.

32. Takeda Y, Jothi R, Birault V, Jetten AM (2012) RORγ directly regulates the circadian expression of clock genes and downstream targets in vivo. Nucleic Acids Res 40(17):8519–8535.

33. Zhao X, et al. (2016) Circadian Amplitude Regulation via FBXW7-targeted REV-ERBα Degradation. Cell 165(7):1644–1657.

34. Bolotin E, et al. (2010) Integrated Approach for the Identification of Human Hepatocyte Nuclear Factor 4α Target Genes Using Protein Binding Microarrays. Hepatol Baltim Md 51(2):642–653.

35. Darsigny M, et al. (2009) Loss of Hepatocyte-Nuclear-Factor-4α Affects Colonic Ion Transport and Causes Chronic Inflammation Resembling Inflammatory Bowel Disease in Mice. PLoS ONE 4(10). doi:10.1371/journal.pone.0007609.

36. Garrison WD, et al. (2006) Hepatocyte Nuclear Factor 4α Is Essential for Embryonic Development of the Mouse Colon. Gastroenterology 130(4):19.e1-19.e.

37. Hayhurst GP, Lee Y-H, Lambert G, Ward JM, Gonzalez FJ (2001) Hepatocyte Nuclear Factor 4α (Nuclear Receptor 2A1) Is Essential for Maintenance of Hepatic Gene Expression and Lipid Homeostasis. Mol Cell Biol 21(4):1393–1403.

38. Li J, Ning G, Duncan SA (2000) Mammalian hepatocyte differentiation requires the transcription factor HNF-4α. Genes Dev 14(4):464–474.

39. Lucas B, et al. (2005) HNF4α reduces proliferation of kidney cells and affects genes deregulated in renal cell carcinoma. Oncogene 24(42):6418–6431.

40. Odom DT, et al. (2004) Control of Pancreas and Liver Gene Expression by HNF Transcription Factors. Science 303(5662):1378–1381.

41. Stoffel M, Duncan SA (1997) The maturity-onset diabetes of the young (MODY1) transcription factor HNF4α regulates expression of genes required for glucose transport and metabolism. Proc Natl Acad Sci 94(24):13209–13214.

42. Zhang EE, et al. (2009) A Genome-wide RNAi Screen for Modifiers of the Circadian Clock in Human Cells. Cell 139(1):199–210.

43. Ramanathan C, et al. (2014) Cell Type-Specific Functions of Period Genes Revealed by Novel Adipocyte and Hepatocyte Circadian Clock Models. PLoS Genet 10(4):e1004244.

44. Michael AK, et al. (2017) Formation of a repressive complex in the mammalian circadian clock is mediated by the secondary pocket of CRY1. Proc Natl Acad Sci 114(7):1560–1565.

45. Xu H, et al. (2015) Cryptochrome 1 regulates the circadian clock through dynamic interactions with the BMAL1 C terminus. Nat Struct Mol Biol 22(6):476–484.

46. Michael AK, et al. (2015) Cancer/Testis Antigen PASD1 Silences the Circadian Clock. Mol Cell 58(5):743–754.

47. Bogan AA, et al. (2000) Analysis of protein dimerization and ligand binding of orphan receptor HNF4α11Edited by M. Yaniv. J Mol Biol 302(4):831–851.

48. Briançon N, Weiss MC (2006) In vivo role of the HNF4α AF-1 activation domain revealed by exon swapping. EMBO J 25(6):1253–1262.

49. Hadzopoulou-Cladaras M, et al. (1997) Functional Domains of the Nuclear Receptor Hepatocyte Nuclear Factor 4. J Biol Chem 272(1):539–550.

50. Sladek FM, Ruse MD, Nepomuceno L, Huang S-M, Stallcup MR (1999) Modulation of Transcriptional Activation and Coactivator Interaction by a Splicing Variation in the F Domain of Nuclear Receptor Hepatocyte Nuclear Factor 4α1. Mol Cell Biol 19(10):6509–6522.

51. Wang J-C, Stafford JM, Granner DK (1998) SRC-1 and GRIP1 Coactivate Transcription with Hepatocyte Nuclear Factor 4. J Biol Chem 273(47):30847–30850.

52. Yoon JC, et al. (2001) Control of hepatic gluconeogenesis through the transcriptional coactivator PGC-1. Nature 413(6852):131–138.

53. Dean S, Tang JI, Seckl JR, Nyirenda MJ (2010) Developmental and tissue-specific regulation of hepatocyte nuclear factor 4-alpha (HNF4-alpha) isoforms in rodents. Gene Expr 14(6):337–344.

54. Tanaka T, et al. (2006) Dysregulated expression of P1 and P2 promoter-driven hepatocyte nuclear factor-4alpha in the pathogenesis of human cancer. J Pathol 208(5):662–672.

55. Torres-Padilla ME, Fougère-Deschatrette C, Weiss MC (2001) Expression of HNF4α isoforms in mouse liver development is regulated by sequential promoter usage and constitutive 3′ end splicing. Mech Dev 109(2):183–193.

56. Takahashi JS (2004) Introduction: Finding new clock components; past and future. J Biol Rhythms 19(5):339–347.

57. Inoue Y, Yu A-M, Inoue J, Gonzalez FJ (2004) Hepatocyte Nuclear Factor 4α Is a Central Regulator of Bile Acid Conjugation. J Biol Chem 279(4):2480–2489.

58. Martinez-Jimenez CP, Kyrmizi I, Cardot P, Gonzalez FJ, Talianidis I (2010) Hepatocyte Nuclear Factor 4α Coordinates a Transcription Factor Network Regulating Hepatic Fatty Acid Metabolism. Mol Cell Biol 30(3):565–577.

59. Stroup D, Chiang JYL (2000) HNF4 and COUP-TFII interact to modulate transcription of the cholesterol 7α-hydroxylase gene (CYP7A1). J Lipid Res 41(1):1–11.

60. Koike N, et al. (2012) Transcriptional Architecture and Chromatin Landscape of the Core Circadian Clock in Mammals. Science 338(6105):349–354.

61. Jung D, Kullak-Ublick GA (2003) Hepatocyte nuclear factor 1α: A key mediator of the effect of bile acids on gene expression. Hepatology 37(3):622–631.

62. Chiang JY, Stroup D (1994) Identification and characterization of a putative bile acid-responsive element in cholesterol 7 alpha-hydroxylase gene promoter. J Biol Chem 269(26):17502–17507.

63. Kamiya A, Inoue Y, Gonzalez FJ (2003) Role of the hepatocyte nuclear factor 4alpha in control of the pregnane X receptor during fetal liver development. Hepatol Baltim Md 37(6):1375–1384.

64. Pizarro A, Hayer K, Lahens NF, Hogenesch JB (2013) CircaDB: a database of mammalian circadian gene expression profiles. Nucleic Acids Res 41(D1):D1009–D1013.

65. Ray A, Prefontaine KE (1994) Physical association and functional antagonism between the p65 subunit of transcription factor NF-kappa B and the glucocorticoid receptor. Proc Natl Acad Sci 91(2):752–756.

66. Yang-Yen H-F, et al. (1990) Transcriptional interference between c-Jun and the glucocorticoid receptor: Mutual inhibition of DNA binding due to direct protein-protein interaction. Cell 62(6):1205–1215.

67. Bosscher KD, Berghe WV, Haegeman G (2006) Cross-talk between nuclear receptors and nuclear factor κB. Oncogene 25(51):6868–6886.

68. Busillo JM, Cidlowski JA (2013) The five Rs of glucocorticoid action during inflammation: ready, reinforce, repress, resolve, and restore. Trends Endocrinol Metab 24(3):109–119.

69. Cain DW, Cidlowski JA (2017) Immune regulation by glucocorticoids. Nat Rev Immunol 17(4):233–247.

70. Caldenhoven E, et al. (1995) Negative cross-talk between RelA and the glucocorticoid receptor: a possible mechanism for the antiinflammatory action of glucocorticoids. Mol Endocrinol 9(4):401–412.

71. Garside H, et al. (2004) Glucocorticoid Ligands Specify Different Interactions with NF-κB by Allosteric Effects on the Glucocorticoid Receptor DNA Binding Domain. J Biol Chem 279(48):50050–50059.

72. Heck S, et al. (1994) A distinct modulating domain in glucocorticoid receptor monomers in the repression of activity of the transcription factor AP-1. EMBO J 13(17):4087–4095.

73. Hatanaka F, et al. (2010) Genome-Wide Profiling of the Core Clock Protein BMAL1 Targets Reveals a Strict Relationship with Metabolism. Mol Cell Biol 30(24):5636–5648.

74. Menet JS, Pescatore S, Rosbash M (2014) CLOCK:BMAL1 is a pioneer-like transcription factor. Genes Dev 28(1):8–13.

75. Rey G, et al. (2011) Genome-Wide and Phase-Specific DNA-Binding Rhythms of BMAL1 Control Circadian Output Functions in Mouse Liver. PLOS Biol 9(2):e1000595.

76. Rudic RD, et al. (2004) BMAL1 and CLOCK, Two Essential Components of the Circadian Clock, Are Involved in Glucose Homeostasis. PLOS Biol 2(11):e377.

